# A simple and efficient *in planta* transformation method based on the active regeneration capacity of plants

**DOI:** 10.1101/2023.01.02.522521

**Authors:** Guoguo Mei, Ao Chen, Yaru Wang, Shuquan Li, Minyi Wu, Xu Liu, Xingliang Hou

## Abstract

Plant genetic transformation strategies serve as essential tools for the genetic engineering and advanced molecular breeding of plants. However, the complicated operational protocol and low efficiency of the current transformation strategies restrict the genetic modification of most plant species. This paper describes the development of a <u>R</u>egenerative <u>A</u>ctivity-dependent *in <u>P</u>lanta* <u>I</u>njection <u>D</u>elivery (RAPID) method based on the active regeneration capacity of plants. In this method, *Agrobacterium tumefaciens* was delivered to plant meristems via injection for inducing transfected renascent tissues. Stable transgenic plants were obtained by subsequent vegetative propagation of the positive renascent tissues. The method was successfully applied for the transformation of plants with strong regeneration capacity, including different genotypes of sweet potato (*Ipomoea batatas*), potato (*Solanum tuberosum*), and bayhops (*I. pes-caprae*). Compared to the traditional transformation methods, RAPID has a markedly high transformation efficiency (up to ~ 40%), shorter duration (less than 4 weeks), and does not require tissue culture procedures. The RAPID method therefore overcomes the limitations of traditional methods for achieving rapid *in planta* transformation, and can be potentially applied to a wide range of plant species that are capable of active regeneration.

## Main

The emerging strategies in biotechnology, including genome editing and high-throughput sequencing, immensely promote the progress of modern agriculture, and are widely employed for improving the agronomic traits of crop plants^1,2^. The accuracy of genetic modification primarily relies on plant transformation, which is difficult to achieve in most plant species^3^. To date, gene editing based on traditional transformation strategies has been successfully realized in only a limited number of representative crops^4^. The functional analysis of genes, exploration of the mechanism underlying important traits, and the genetic improvement of potentially valuable plant species face several challenges owing to the lack of a universal transformation technology.

The existing methods for the genetic transformation of plants primarily include particle bombardment, viral vector delivery, and delivery of *Agrobacterium* sp. for transferring synthetic exogenous information into genomes. Of these, the *A*. sp.- mediated transformation of plants is mostly employed owing to its high efficiency and reliability^5–7^. Following inflorescence infection, *A. tumefaciens* directly contacts germ cells of the model plant, *Arabidopsis thaliana,* to induce the formation of the transformed progeny^8,9^. Although the floral dip strategy is ideal for plant transformation, the method is unsuitable for other flowering plants. Most plant transformation methods involve the conversion of *A*. sp.-infected dedifferentiated calli into regenerated plantlets through tissue culture^5^. However, this strategy is primarily limited by numerous factors, including the plant species, culture conditions, several genetic and technical instabilities^10^, expensive and complicated processes for culturing transformed cells, high time consumption^11^, and low transformation efficiency.

Several advances have been made in plant genetic transformation technologies in recent years, based on tissue culture method or nonsterile conditions. In one strategy, RNA viruses and mobile elements were used to guide dsRNA components into plant meristems. This method elevates the efficiency of heritable gene editing but possibly requires a tissue culture procedure for generating the primary Cas9 transgenic plant^12^. In another strategy, meristem identity regulators were used to promote the regeneration of transformed tissues without sterile operation procedures^13^. In a recent study, *A. rhizogenes* was used for directly inducing the regeneration of transgenic organs^14^. Additionally, numerous studies have employed various delivery intermediaries, including viruses and nanomaterials for plant transformation^6,15^. These transformation strategies offer different solutions for improving the transformation efficiency of plants. However, it is necessary to develop novel transformation strategies that can be easily and universally applied in various plant species for overcoming the technological limitations and challenges arising from plant diversity.

This paper describes an efficient, easy-to-use *A. tumefaciens*-mediated transformation system, denoted as <u>R</u>egenerative <u>A</u>ctivity-<u>d</u>ependent *in <u>P</u>lanta* <u>I</u>njection <u>D</u>elivery (RAPID), which has the advantages of a high transformation rate, short duration, user-friendly operational protocol, and no requirement for tissue culture. The study confirmed that the RAPID method significantly promoted the genetic transformation of plants with strong regeneration capacities, including sweet potatoes, potatoes, and bayhops. The RAPID strategy can therefore serve as a potentially powerful tool for the genetic modification of plants that propagate vegetatively.

## Results

### Implementation of a simple and effective *in planta* transformation delivery system in sweet potato

Owing to the natural regeneration capacity, plants can be propagated from excised organs, including leaves, stems, and roots^16–18^. It is therefore possible to obtain independent transformants by *in planta* regeneration instead of tissue culture. Sweet potato (*Ipomoea batatas* L.), a well-known tuberous crop with strong vegetative propagation capacity in the stems and roots^19^, was selected in this study for developing the tissue culture-free transformation approach. Various methods, including soaking, vacuum infiltration, injecting, and other strategies, were tested for delivering *A. tumefaciens* into various plant tissues, including the leaves, stems, flowers, and roots. The transformation efficacy of various combinations of post-transfection culture substrates, including water, nutrient solution, sand, and solid media, was also evaluated. The findings revealed that the direct injection of stem segments with a subsequent culture in the soil substrate achieved an effective and stable transformation (**Fig. 1**).

**Fig. 1.**
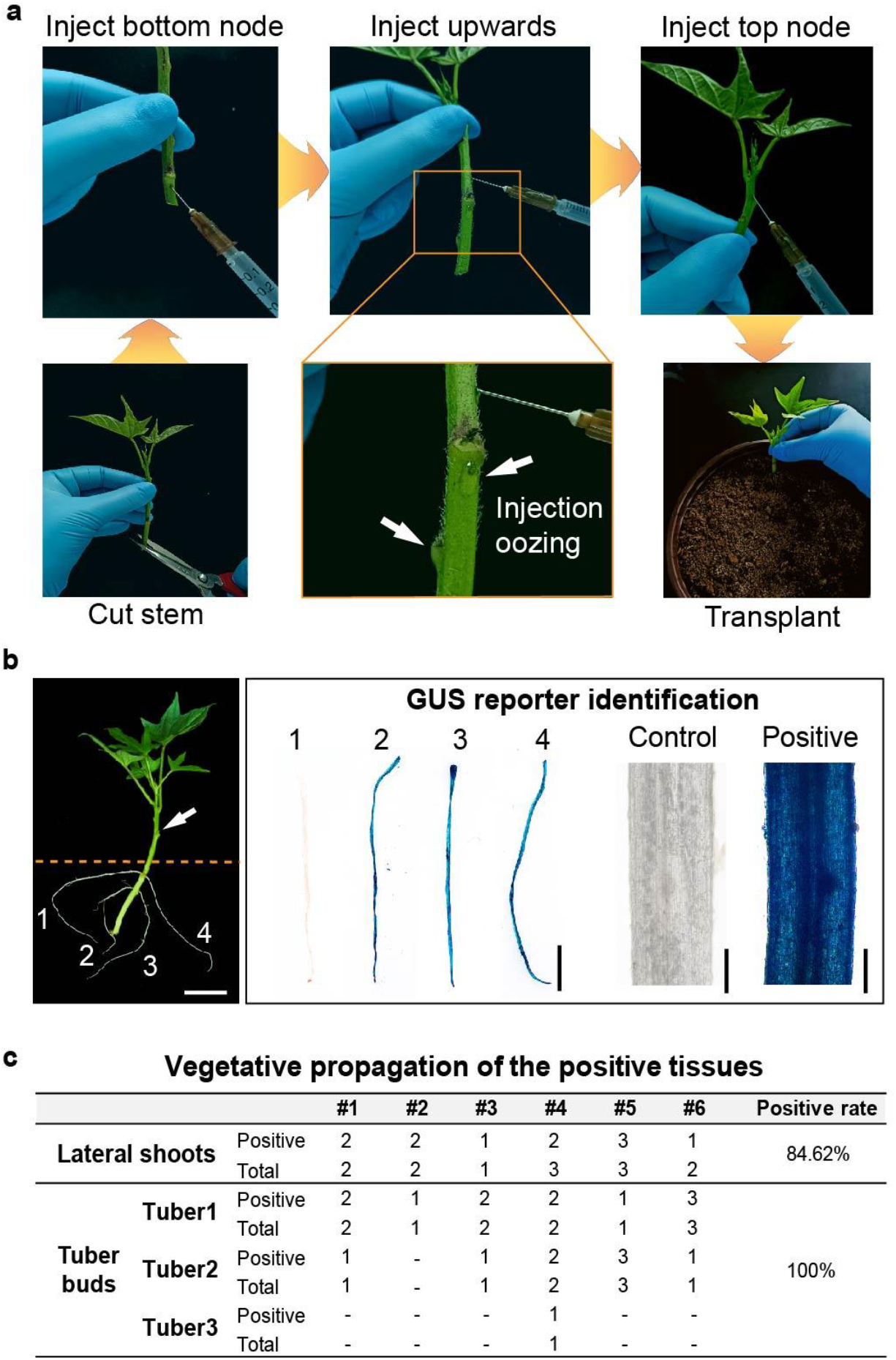
Operating procedure of the stem-injection delivery system. **a**, The operating procedure of the stem-injection delivery system. The healthy stems of sweet potato plants with several nodes were excised, and each node was injected upwards until the solution oozed from the adjacent pinholes and excised end. The injected stems were planted into the soil substrate. **b**, Evaluation of the transformation of renascent roots. The adventitious roots sprouted spontaneously under the soil within 1 week (below the yellow line) and were selected for GUS staining. Scale bar, 1 cm (left), 0.5 cm (middle), 0.5 mm (right). **c**, Vegetative propagation of the positive tissues. Independent transgenic plants were further obtained by the vegetative propagation of the positive lateral shoots, or from the buds that sprouted from the positive tubers. Positive rate = average (positive/total) × 100%.

The protocol of the stem-injection delivery strategy is depicted in **Fig. 1a**. Healthy sweet potato stems bearing several nodes were excised, and each node was injected upward until the injection liquid oozed from the other pinholes and excised ends. The injected stems were subsequently planted into the soil substrate (**Fig. 1a**), and adventitious roots spontaneously sprouted within 1 week after soil cutting (**Fig. 1b**). The positive transformants were rapidly detected from the adventitious roots. *GUS* (*β-glucuronidase*) was selected as the reporter gene for evaluating the transformation of the renascent roots (**Fig. 1b**). Continuous cultivation of the stem cuttings generated transgenic renascent leaves, lateral buds, and tubers from the adventitious roots, and the positive signal was detected in these tissues **(Supplementary Fig. 1)**. The independent transgenic plants were obtained by subsequent vegetative propagation of the positive lateral shoots within a short term, or from the buds that sprouted from the positive tubers (**Fig. 1c**). The study therefore preliminarily established an effective *in planta* transformation method by taking advantage of the active regeneration capacity of sweet potato plants.

### Optimization of the *in planta* transformation system

The offspring that were vegetatively propagated from the positive renascent shoots and tubers were genotyped, and the results demonstrated that the examined plant tissues were also positive (**Fig. 1c**). The finding suggested that the transgenic events produced by the *in planta* method are stable and have a low chimeric rate. We subsequently aimed to improve the transformation efficiency of the stem injection method. To this end, we first screened different strains of *A.* sp. commonly used for plant transformation, including the AGL1, GV3101, EHA105, and LBA4404 strains of *A. tumefaciens*, and the K599 strain of *A. rhizogenes*^9,20,21^. Following transfection, the renascent adventitious roots were selected for GUS staining, and the positive rate (ratio of positive roots per positive plant × ratio of positive plants in all the injected plants) was determined for evaluating the transformation efficiency (**Supplementary Table 1**). The results demonstrated that the AGL1 strain of *A. tumefaciens* had the highest transformation efficiency (28%), followed by the GV3101 and EHA105 strains (19%), while transformation with the LBA4404 strain did not generate any positive roots. The K599 strain of *A. rhizogenes* had a weak transformation efficiency of < 2% (**Fig. 2a**). Therefore, the AGL1 strain of *A. tumefaciens* was found to be most suitable for the transformation of sweet potato using this system.

**Fig. 2.**
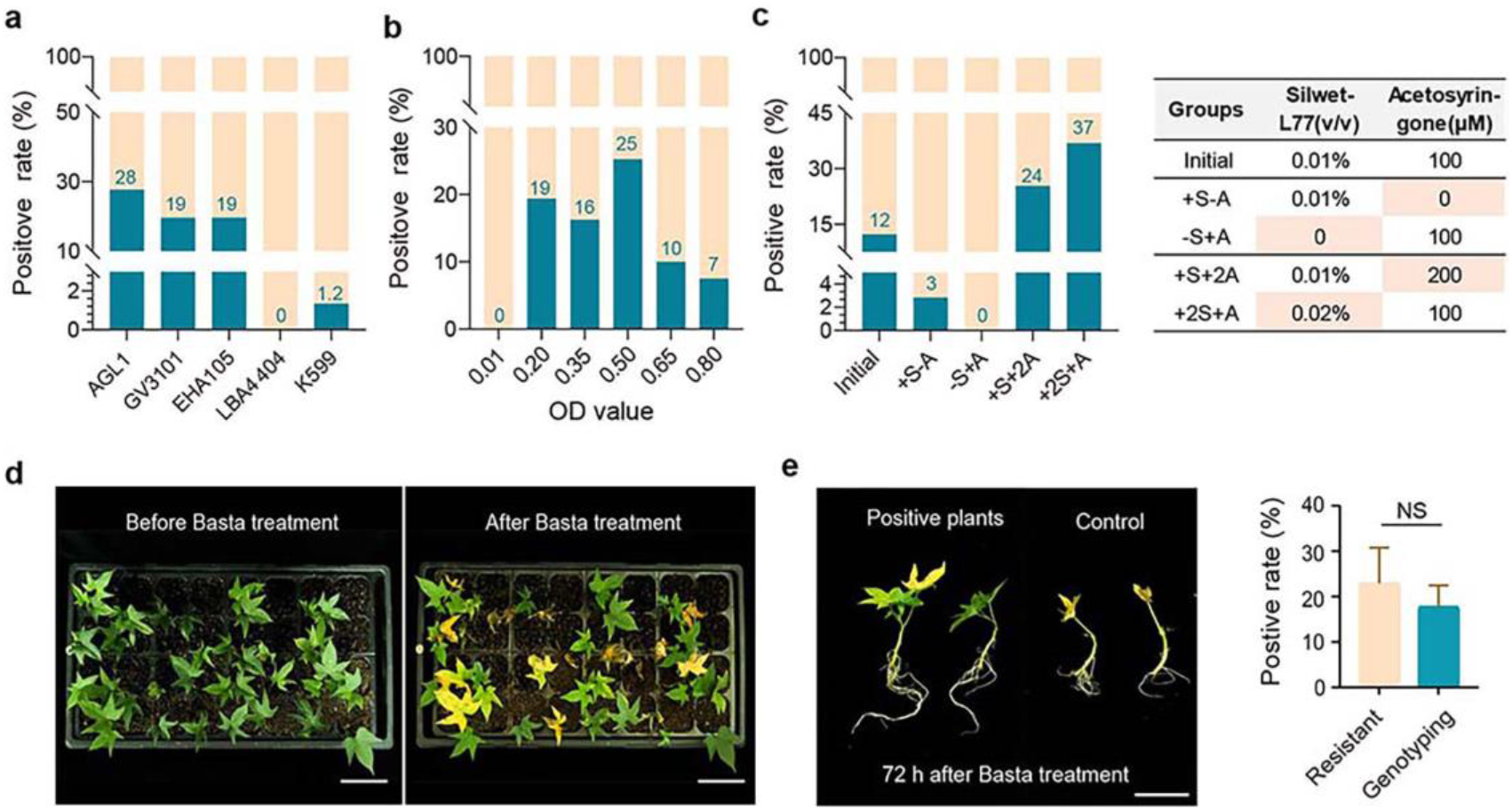
Optimization of the stem-injection delivery system. **a**, Transformation efficiency of different *A*. sp. strains. **b**, Transformation efficiency of *A*. sp. strains at different OD values. **c**, Transformation efficiency following the addition of different additives. Positive rate = ratio of the number of positive roots per positive plant × ratio of the number of positive plants among all injected plants. **d**, Bulk selection of transgenic materials based on phosphinothricin (Basta) resistance. The phenotype of the injected plants before and after spraying Basta. **e**, Comparison of the morphology of the positive plants and wild-type (WT) after 72 h of spraying Basta. The bar chart shows the positive rate of Basta resistance and genotyping of the screened plants. Positive rate = number of positive plants/total number of plants (%). The data are presented as the mean ± SD of 10 biological replicates (two-tailed Student’s *t*-test, NS, no significance, P > 0.05). Scale bar, 5 cm in **d**, 5 cm in **e**.

The optimal optical density (OD), which is another important variable for *A.*sp.-mediated transformation, was subsequently determined. The AGL1 strain was cultured and diluted to generate a series of concentrations with OD values ranging from 0.01 to 0.80, and the cultures were subsequently injected into sweet potato stems for investigating the GUS-positive roots. The findings revealed that OD_0.5_ was the optimal OD for transformation (**Fig. 2b**, **Supplementary Table 1**). Previous studies reported that chemical additives, including Silwet-L77 and acetosyringone, can significantly promote the transformation rates of the floral dip method for the transformation of *Arabidopsis* and the tissue culture-based transformation methods in some plants^8,22^. The initial infecting solution for injection, containing 0.01% Silwet-L77 and 100 μM acetosyringone, was prepared as previously described (**Fig. 2c**)^8^. Different combinations of these two additives were compared, and the results demonstrated that the transformation failed in the absence of Silwet-L77, which indicated that the surfactant component, Silwet-L77 (S), is critical for successful transformation. Acetosyringone (A) also had a prominent effect in promoting the transformation efficiency. Notably, the findings revealed that the transformation efficiency was significantly improved to 37% when 0.02% Silwet-L77 and 100 μM acetosyringone (2S+A) were added to the solution (**Fig. 2c**).

To improve the screening throughput, we attempted to select multiple explants using herbicides and antibiotics (**Fig. 2d, Supplementary Fig. 2**). The successful transformants carrying resistance genes can resist the damages caused by externally applied compounds, and these positive plants were further confirmed by genotyping (**Fig. 2e, Supplementary Fig. 2**).

The present study describes the development of a novel method for plant transformation that does not require a sterile operation strategy. The transformation rate of the method was subsequently elevated to nearly 40% and the period was shortened to only 1 month via multiple optimization steps. The method described herein exhibited remarkable superiority over the traditional tissue culture method used for sweet potato (**Table 1**). The transformation method developed in this study was denoted as the Regenerative Activity-dependent *in Planta* Injection Delivery (RAPID) method.

**Table 1.**
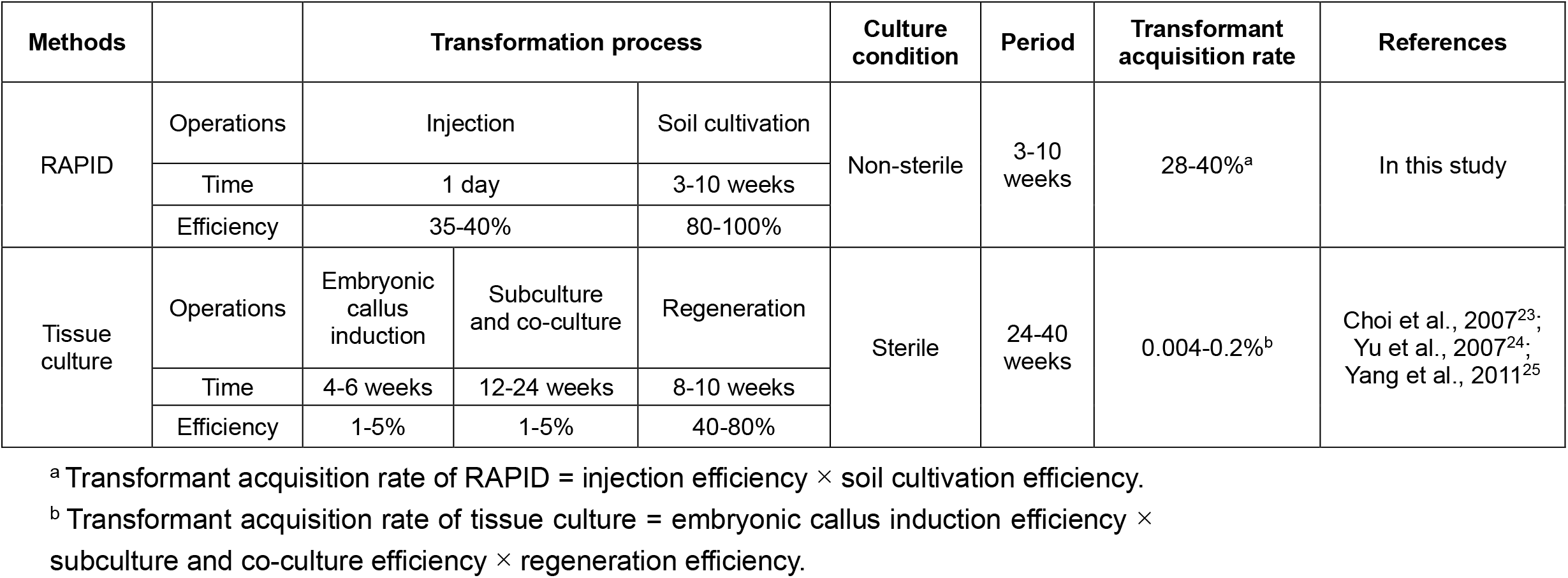
Comparison of RAPID and traditional tissue culture method in sweet potato.

### The RAPID method can deliver multiple reporter vectors and is a reliable gene editing tool

In order to verify the applicability of the RAPID system, diverse reporter genes were selected for verifying the transformation. The application of fluorescent reporters can aid in examining the transformation of plants in a living state; however, the interference of spontaneous biological fluorescence needs to be considered as well^23^. We determined the spontaneous fluorescence spectra of sweet potato tissues using a confocal microscope (**Supplementary Fig. 3**). In order to evade the interference due to spontaneous biological fluorescence, the mScarlet red fluorescent protein (the excitation and emission wavelengths were at 569 nm and 593 nm, respectively) was selected as the transformation reporter. As expected, most of the positive renascent tissues identified by polymerase chain reaction (PCR) exhibited an obvious red fluorescence. Furthermore, the strong fluorescence signals were maintained in the leaves of the individual transgenic plants regenerated from the transformants (**Fig. 3a**), suggesting that mScarlet is a suitable fluorescence reporter for studying the transformation of sweet potato. The RUBY reporter system was subsequently tested in this study. RUBY can produce visible red accumulation in living plants, and is also used for monitoring transformation events under non-invasive conditions^24^. The RUBY transformants in our delivery system were similar to those reported in other species^24,25^, and exhibited an obvious red phenotype in the positive sweet potato plants (**Fig. 3b**).

**Fig. 3.**
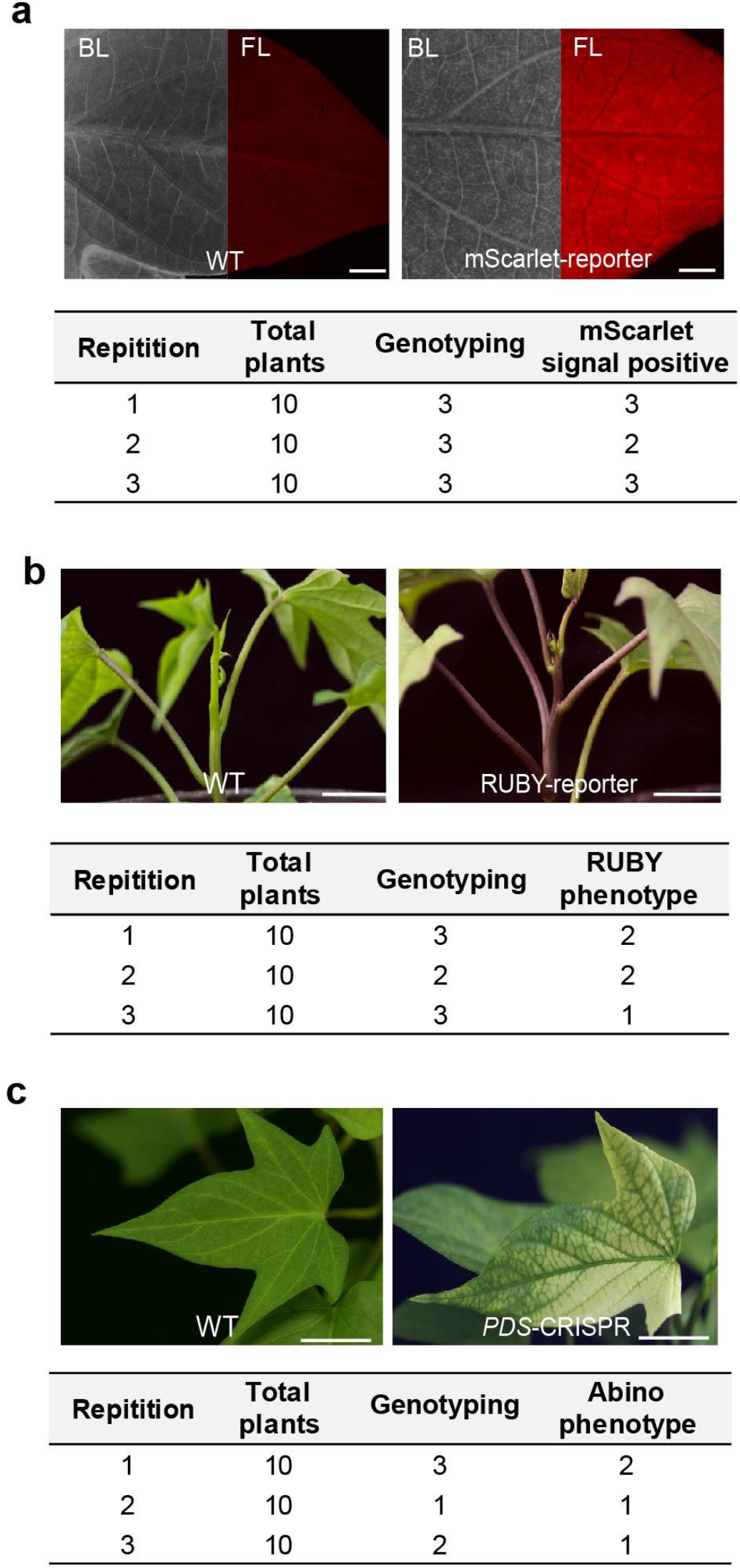
Transformation applicability of reporter vectors and gene editing tools in RAPID. **a**, Applicability of the mScarlet reporter in transformation. Scale bar, 0.5 cm. The table depicts the statistics of the transformation results obtained from three independent replicates. BL, bright light; FL fluorescent light. **b**, The applicability of the RUBY reporter in transformation. Scale bar, 2 cm. The table depicts the statistics of the transformation results obtained from three independent replicates. **c**, The applicability of the CRISPR-Cas9 tool in transformation following *PDS* knockout. Scale bar, 2 cm. The statistics of the transformation results obtained from three independent replicates are depicted in the table.

Gene editing systems serve as important tools for plant genetic research. To verify whether the RAPID system can be compatible with gene editing tools, the *phytoene desaturase* (*PDS*) homolog gene (g31261) of sweet potato was knocked out by CRISPR-Cas9^26^. The loss of *PDS* function produces distinct albino phenotypes in different plants^27^, and the transgenic renascent shoots gradually developed an obvious albino phenotype in this study (**Fig. 3c**). Taken together, the results suggested that the RAPID method is reliable for the delivery of diverse report vectors and application in studies on gene editing.

### RAPID produces positive renascent tissues by inducing the successful transfection of meristematic cells

RAPID is an efficient method for plant transformation, and has a high transformation efficiency; however, certain transient transfection methods, including virus-induced gene silencing (VIGS) and leaf-agroinfiltration, can also produce a high transformation rate^28^. The key difference between these methods and RAPID is that the latter can generate stable regenerative transgenic plants. Lateral buds and adventitious roots are known to develop from the meristematic cells of phloem tissues^17,18,29^; we therefore speculated that the RAPID method might induce the transfection of tissues with regeneration capacity. In order to test this hypothesis, mScarlet was selected as the reporter gene for analyzing the stem biopsies following transfection. The untreated plants in the control group exhibited a spontaneous fluorescence in the epidermal layer of the stems. Interestingly, the transformants exhibited obvious fluorescence signals in the interior cross-section of the stem (**Fig 4a**). Further observations revealed that these signals were primarily localized in the meristematic regions of the phloem, including the cambium and endodermis, which are involved in the differentiation of lateral buds and adventitious roots in sweet potato (**Fig 4b**)^19,30^. Consistently, the lateral buds and adventitious roots exhibited strong positive signals (**Supplementary Fig. 4**), and regenerated transgenic offspring. These observations suggested that the RAPID method can induce the transfection of the meristematic zone via the phloem *in planta*, and generate stable regenerated plants from the lateral and tuber buds (**Fig. 4c**). The period between injection and the generation of individual transgenic plants from the lateral buds and tuber buds was 3–4 weeks and 6–8 weeks, respectively. All the positive lines obtained from the tuber buds were non-chimeric and supported the notion that the transgene originated from one or a few meristematic cells.

**Fig. 4.**
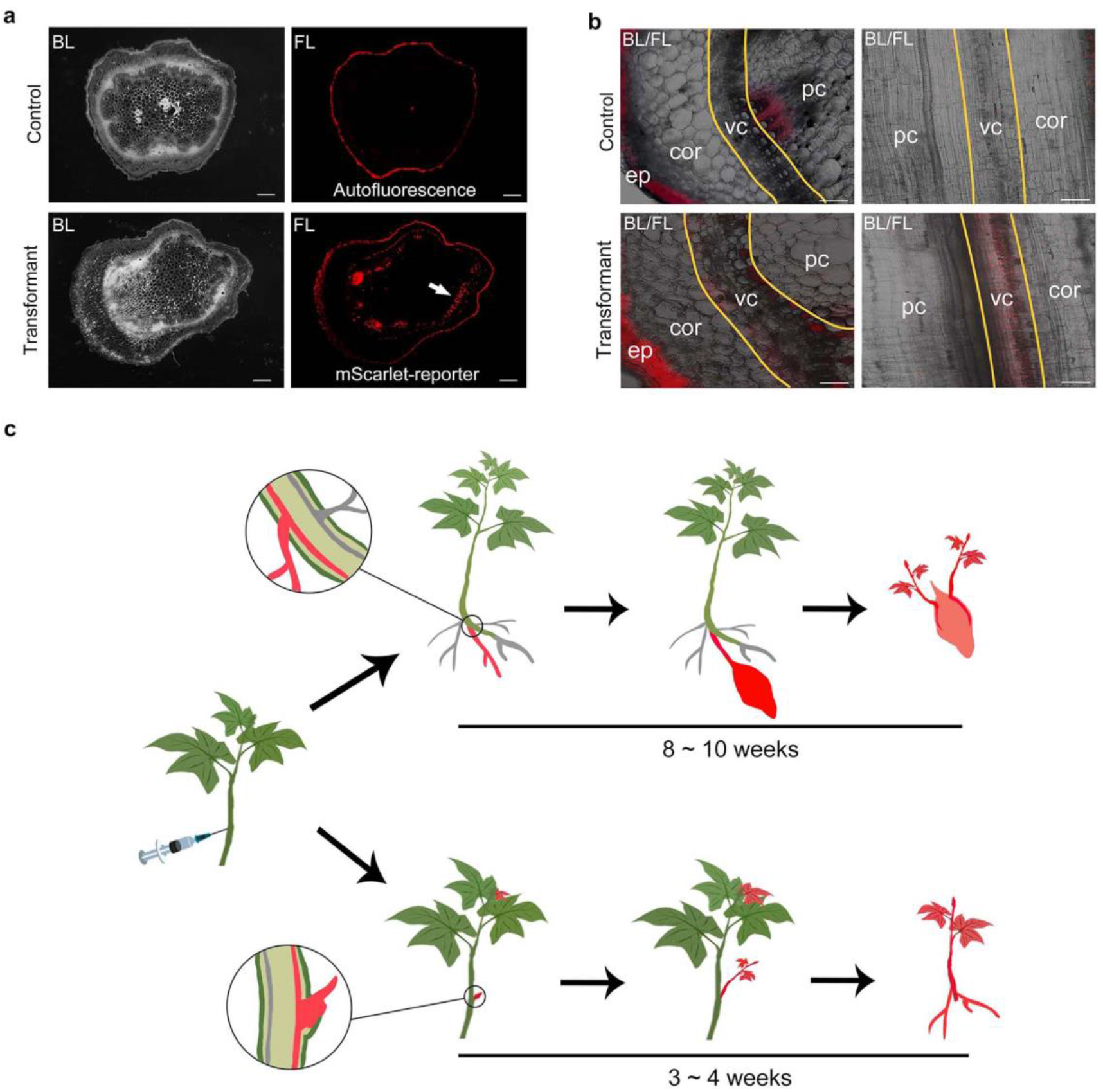
*A.* sp. was directly delivered to the phloem in the RAPID system. **a**, Fluorescence due to mScarlet in the cross-section of sweet potato stems. The white arrow indicates the obvious fluorescence signal in the transformant tissue. Scale bar, 0.5 mm; bright light, BL; fluorescent light, FL. **b**, Histological observation of the signal due to the mScarlet reporter. The area within the yellow lines in the transverse (left) and longitudinal sections (right) of the stems depict the signal due to mScarlet in the transformant; scale bar, 0.1 mm. Vascular cell, vc; parenchymal cell, pc; cortex, cor; epidermis, ep. **c**, Working model of the acquired transformed generations following direct transfection with the RAPID system.

### RAPID is applicable to different plant species

In order to evaluate the applicability of the RAPID method to different genetic backgrounds, we first tested its efficacy in different varieties of sweet potato, including the major cultivars (Guangshu87, Pushu32, Longshu9, and Jishu26), a cultivar with high anthocyanin content (Guangzishu8), and a genome-sequenced cultivar (Taizhong6)^31^ (**Fig. 5a**). To detect the transgenic materials on a large scale, the artificial light source method was employed for detecting the GFP signals^32^. The stems were peeled to nullify the interference due to the spontaneous biological fluorescence of the epidermis. Strong green fluorescence was observed in the peeled stems of the GFP transformants, suggesting that the RAPID method can be applied for the transformation of sweet potato plants with different genetic backgrounds. Additionally, the transformation rate of the RAPID method was high in sweet potato (12.5–37.5%; **Fig. 5a**).

**Fig. 5.**
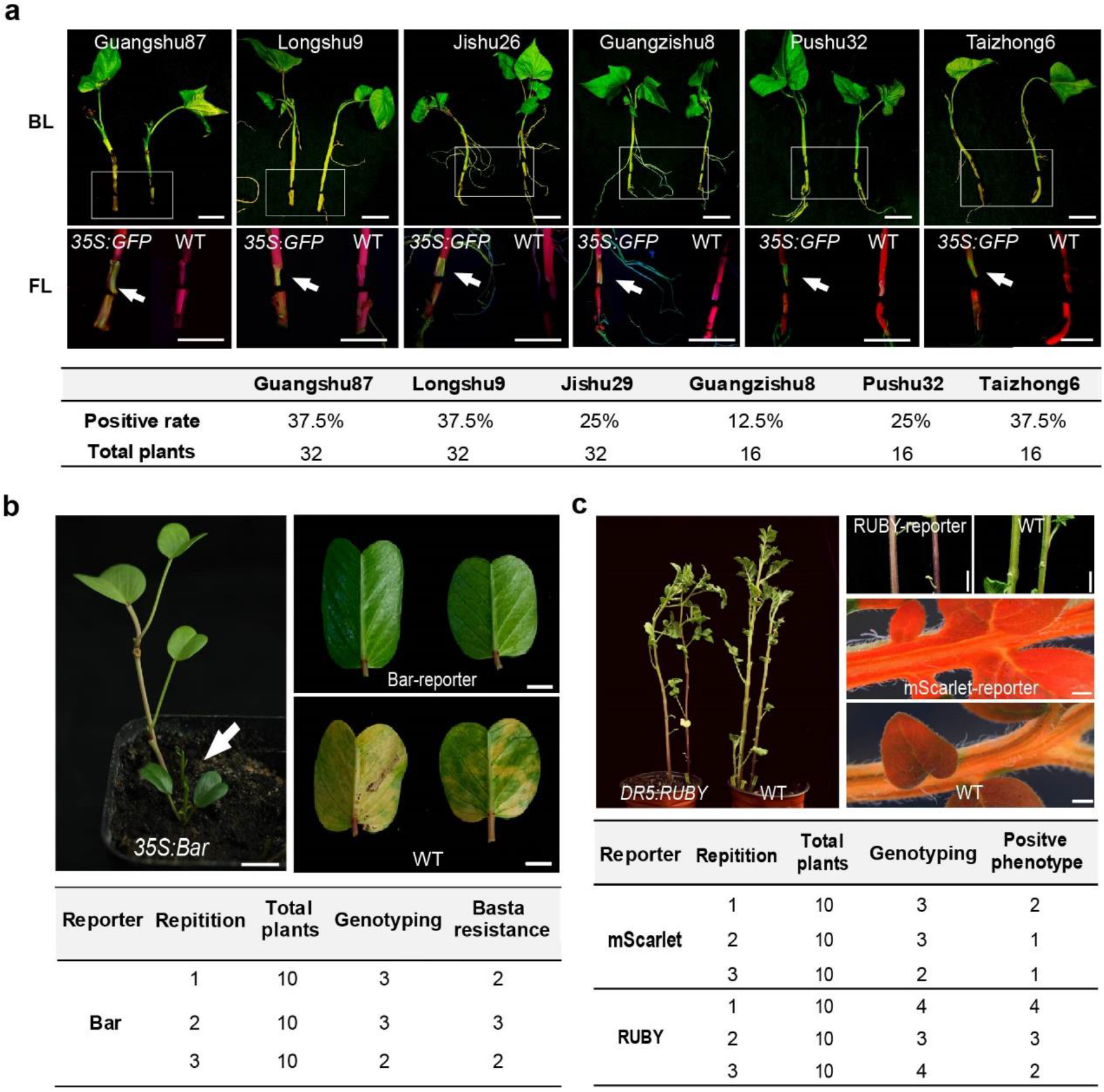
Application of the RAPID system in other plant species. **a**, Transformation of different varieties of sweet potato using the RAPID method. The stem was peeled and observed under light and dark conditions (within the box). The arrow indicates the green fluorescence due to the GFP reporter (*35S:GFP*). Positive rate = number of positive plants/total number of plants (%). Scale bar, 2 cm; bright light, BL; fluorescent light, FL. **b**, Transformation of bayhops. The white arrow indicates the renascent shoots from the transformed stems. The leaves from the transgenic shoots carrying the *Bar* gene (*35S:Bar*) retained the green fluorescence under phosphinothricin (Basta) treatment compared to the WT. Scale bar, 2 cm (left), 1 cm (right). The table depicts the statistics of transformation results obtained from three independent replicates. **c**, Transformation of potato. The transgenic materials carrying the RUBY (*DR5:RUBY*) and mScarlet (*35S: mScarlet*) reporters exhibited positive phenotypes. Scale bar, 2 cm (upper), 0.5 cm (lower). The table depicts the two statistics of transformation results obtained from three independent replicates.

The present study also determined whether the RAPID method can be applied to other plant species. Bayhops (*I. pes-caprae* L.) is a coastal plant with strong adaptation to barren and saline conditions, and has a high potential for application in vegetation repair. In this study, bayhops was selected for analyzing whether the RAPID method can be applied for the transformation of this plant species. Bayhops and sweet potato belong to the Convolvulaceae family; however, unlike sweet potato, the vegetative propagation of bayhops occurs via stolons rather than via storage roots. There are no established transformation systems for bayhops to date. In this study, the stems of bayhops plants were excised and transformed using the RAPID system by transferring the *Bar* gene. PCR detection revealed that out of the 30 plants transfected in this study, the renascent shoots of 8 plants were positive, and most of the regenerated transgenic plants that developed from these shoots exhibited herbicide resistance (**Fig. 5b**). Taken together, the findings suggested that bayhops can be efficiently transformed using the RAPID method.

Potato (*Solanum tuberosum*. L) is the fourth largest food crop in the world and it exhibits a strong ability of vegetative propagation via tubers^33^. This feature enables the application of the RAPID method for the transformation of potato plants. In order to determine the transformation efficacy of the RAPID method in potato, the detached stems of potato plants were initially infected by injections and cultured in the soil. However, the efficacy could not be precisely determined owing to the unstable survival rate of the stem cuttings of potato. The tubers were subsequently selected as the transformation receptor owing to their stronger natural regeneration capacity compared to stems. We injected *A.* sp. mixed with exogenous growth regulators of 6-BA (6-benzyladenine) and NAA (naphthylacetic acid) into the skin of the tuber around the buds. The growth regulators induced sprouting without causing any undesirable genetic alterations (**Supplementary Fig. 5**). Renascent shoots emerged from the tubers after 1–2 weeks, and the transformants exhibited strong phenotypes of mScarlet and RUBY in the leaves and stems (**Fig. 5c**). The transformation rate was nearly 40%, which was markedly higher than that of the traditional transformation method for potato (<1%)^34^.

Taken together, the above findings revealed that by using selectively targeted tissues and optimized conditions, the RAPID method can be widely applied for the genetic transformation of various plant species to generate transformants with active regeneration capacity.

## Discussion

The existing methods for plant transformation commonly deliver exogenous genes into plants by infecting with *A.* sp., and the majority of these strategies rely on tissue culture procedures^35^. However, the methods for plant transformation often have low transformation efficiency and are highly time-consuming owing to the limitations of the culture conditions, challenges in different plant species, and issues during technical operation, which poses as bottlenecks in plant genetic research and application efforts. In this study, we developed a simple *in planta* transformation strategy, denoted as RAPID, that delivers genes by directly injecting *A. tumefaciens* into plants. The RAPID method overcomes some of the main limitations of the current transformation strategies as described hereafter. First, the RAPID method does not require tissue culture, special induction strategies, or treatments that are necessary in the existing plant transformation methods^12,13^. Successful transformation with the RAPID method only relies on the active regeneration capacity of plants, which substantially increases the transformation efficiency, shortens the duration, and reduces the possibility of genetic mutations caused by cellular dedifferentiation and redifferentiation. Furthermore, RAPID is compatible with different genetic modification strategies, including ectopic expression or gene editing, and with different strains of *A.* sp.; therefore, transformation with RAPID is relatively simple, flexible, and can be employed for diverse purposes. For instance, *A. rhizogenes* has been used for achieving plant transformation under non-sterile conditions. Several recent transformation strategies have also employed *A. rhizogenes* for inducing the regeneration of plant organs^14,36^. However, *A. rhizogenes* potentially causes persistent abnormal plant growth owing to the presence of *rol* genes that are randomly inserted into the genome^37–39^. RAPID can use *A. tumefaciens* as the mediator of infection, and the issues with *A. rhizogenes* can therefore be evaded. Additionally, there is no limitation of specific explants, and the method has a widespread regeneration capacity across different plant species. Theoretically, RAPID can be successfully applied to other plants, besides those tested in this study.

The floral dip method is typically used for the transformation of plants such as *Arabidopsis*, in which the transfected germ cells act as precursors that pass the transgenic information to seeds to generate transgenic plants^8^. RAPID is similar to the floral dip method in that the meristematic cells are transfected via direct contact with *A.* sp., which subsequently leads to the regeneration of positive renascent tissues and transgenic offspring via vegetative propagation. It has been reported that bacteria and viruses can spread through the phloem channels after entering the cortex^40^, which enables direct contact between meristematic cells and *A*. sp. (**Fig. 4**). The RAPID system delivers *A.* sp. into the phloem to ensure that the core cells can be fully transfected *in planta*, and may be one of the reasons underlying the high transformation rate of this method. This hypothesis is also supported by the fact that the Silwet-L77 surfactant is critical for successful transformation using RAPID (**Fig. 2c**). However, the chimeric rate of the regenerated tissues is very low owing to asexual succession (**Fig. 1c**). The germ cells of the secondary plants may carry the genetic information and possibly allow the isolation of genetic information via sexual reproduction.

The strong vegetative propagation ability of plants is the key for obtaining regenerated transgenic plants with the RAPID method. Vegetative propagation is achieved via meristematic cells that actively divide by mitosis and allow the widespread and rapid primary growth of plants, which results in the production of specialized, permanent plant tissue systems for regeneration^41^. These findings indicate that RAPID can be theoretically applied to all plant species that develop into independent plants via vegetative propagation. The chimeric rate is potentially reduced by the regeneration from positive adventitious organs. Because the tubers of sweet potato develop from adventitious roots originating from one or a few meristematic cells in the cortex, all the transgenic plants that regenerated from the tubers were positive in this study. However, further studies are necessary for confirming whether this also occurs in other plant species.

In conclusion, a simple and efficient *A.* sp.*-*mediated transformation system was developed, which overcomes the technical limitations of the existing transformation strategies. The potential wide application of the method for the transformation of different plant species can be achieved via the specific optimization of the strain of *A.*sp. used, tissues infected, culture conditions, and other factors.

## Methods

### Plant materials and growth conditions

The materials of sweet potato (*I. batatas* L.), potato (*S. tuberosum* L.), and bayhops (*I. pes-caprae* L.) used in this study were grown under normal conditions (28°C, 10-h/14-h light/dark cycle, and light intensity of ~200 μmol m^-2^ s^-1^).

### Vector construction

The cloned DNA was amplified and assembled using the Phanta Max Super-Fidelity DNA Polymerase (P505, Vazyme, China) and a ClonExpress II One Step Cloning Kit (C112, Vazyme, China). pCambia1301 was used for the GUS reporter vector (*35S:GUS*). The coding region of *GUS* contained a modified *Catalase* intron from castor bean to ensure that the gene was only expressed under the *35S* promoter of the transformed plant and not expressed in *A*. sp. The coding sequence (CDS) of *mScarlet* was cloned into the pTF101 plasmid^42^ for generating the *35S:mScarlet* reporter vector. The GFP reporter was expressed under the *35S* promoter using the *35S:mGFP* plant binary plasmid, as previously described^43^. The RUBY reporter vector was supported by the *DR5:RUBY* plant binary plasmid, as previously described^24^. The *PDS*-CRISPR construct was generated using the CRISPR-Cas9 system, as previously described^44^. The synthetic guide RNA (sgRNA) targeted the *Ipomoea batatas phytoene desaturase* gene (*IbPDS;* g31261, https://sweetpotao.com/gRNAdesigner) was inserted into the CRISPR-Cas9 binary vector, which was driven by the *AtU6* promoter, while *Cas9* was driven by the *35S* promoter. The sequences of the corresponding vectors are available at https://www.addgene.org, and the template and primers used for vector construction are listed in **Supplementary Table 2**.

### RAPID system

#### *A.* sp. culture and activation

The AGL1, GV3101, EHA105, and LBA4404 strains of *A. tumefaciens*, and the K599 strain of *A. rhizogenes*^45^ carrying the target plasmids were plated on Luria-Bertani (LB) solid medium supplemented with the corresponding antibiotics, and cultured from a single bacterial colony at 28°C for 2–3 days. The freshly grown *A*. sp. was gently washed off from the medium with a wash buffer (10 mM MgCl_2_ and 100 μM acetylsyringone; pH 5.6) and centrifuged at 3000 g for 5 min for collecting the bacteria. The collected *A*. sp. was subsequently diluted to an OD_600_ of 0.5 with an infiltration buffer (1/4 Murashige & Skoog (MS), 0.1 % sucrose (w/v), 100 μM acetylsyringone, and 0.02% Silwet L-77 (v/v); pH 5.6), and prepared for subsequent analyses.

#### Transformation of sweet potato

Stem segments, approximately 10 cm long, and bearing 3–4 nodes and 2–3 mature leaves, were excised from healthy sweet potato plants. The infiltration buffer containing the activated *A*. sp. was drawn into a 1-mL syringe and injected into the lower excised end of the stem segment until the solution oozed from the upper end. The infiltration buffer was subsequently injected upwards into each node with the syringe until the solution oozed from the adjacent pinholes, to ensure that the infection solution was completely distributed in the stem segments. The injected stem segments were transplanted in sandy soil and cultured in a growth chamber for 1–2 days under dark conditions, and subsequently returned to normal conditions under a light cycle (28°C, 10-h/14-h light/dark cycle, and light intensity of ~200 μmol m^-2^ s^-1^). The nascent shoots and adventitious roots derived from the stem cuttings after 2–4 weeks were selected for identifying the positive transformants by molecular detection studies and phenotypic selection. The positive tubers were harvested after 8–10 weeks and sprouted for obtaining the independent transgenic lines by vegetative propagation.

#### Transformation of bayhops

The process of transformation of bayhops was similar to that used for sweet potatoes. Approximately 10 cm-long stem segments bearing several nodes and mature leaves were excised and transplanted in sandy soil following injection with *A*. sp. using a 1-mL syringe. The nascent shoots were obtained after 2–3 weeks and used for determining the positive shoots and phenotypic selection. The independent transformants were obtained by vegetative propagation.

#### Transformation of potato

The previously described method of pretreatment with *A*. sp. was used for the transformation of potato plants; however, the growth regulators NAA (2.5 mg/L) and 6-BA (0.5 mg/L) were added to the infiltration buffer used for the transformation of potato. The fresh tubers of potato were cut, and holes were pricked in the epidermis with the needle of a 1-mL syringe. The infiltration buffer was aspirated with a needleless syringe and the buffer was injected beneath the skin of the tubers, close to the regions around the buds. The injected tuber segments were transplanted in sandy soil. The new buds were unearthed after 1–2 weeks for the detection of positive tubers and phenotypic analyses. The positive tubers were harvested after 6–8 weeks and propagated for developing independent transgenic lines by vegetative propagation.

#### Selection of transformants

Phosphinothricin (Basta^®^, 0.002%, v/v, CB2471, Coolaber, China) and hygromycin (50 μg/mL, H370, XMJ Scientific, China) were sprayed and applied to the aboveground parts for resistance screening. The treated plants were selected after 72 h, and the leaves of the non-transformed plants appeared wilted and yellow, while the positive plants were less affected.

#### GUS staining

The GUS staining assay was performed as previously described^46^. The adventitious roots were immersed in the work solution (50 mM Na_2_PO_4_, 0.5 mM K_3_Fe(CN)_6_, 0.5 mM K_4_Fe(CN)_6_, and 2 mM X-Gluc; pH 7.0) under vacuum for 5 min and subsequently allowed to react at 37°C for 4–6 h. The immersed tissues were placed in a decolorization solution (70% ethanol and 30% acetic acid) for 2-3 times for decolorizing the tissues. The decolorized samples were biopsied and observed under a stereoscope (M165C, Leica, Germany).

#### Fluorescence detection

The fluorescence of the mScarlet reporter in the live transgenic tissues was observed under a fluorescent stereoscope (M205 FA, Leica, Germany) with a red fluorescence filter (Exc 540–580 nm, Em 593–667 nm). The spontaneous fluorescence spectra of sweet potato were observed under a confocal microscope (SP8 STED 3X, Leica, Germany). One-week-old adventitious roots were freshly selected for testing the root samples, and the third mature leaf of the nascent shoots was selected for testing the leaf samples. The materials of transgenic sweet potato containing the GFP reporter were observed using the dual-wavelength fluorescent protein excitation light source (Exc 440 nm, Em 500 nm, 3415RG, LUYOR, China). The regions near the phloem at the base of the stems were peeled for nullifying the spontaneous biological fluorescence of the epidermis, and the whole plant was irradiated by a light source under dark conditions. Green fluorescence was detected in the inner region of the stems of positively transformed plants.

#### Statistical analyses

To evaluate the transformation efficacy, at least 10 biological replicates were performed, and the numbers of replicates are depicted in the figures. The positive rate was calculated using the equation: positive rate = [average (positive roots/total number of roots)] × positive strains/total number of strains × 100%. For molecular biology analysis, at least three individual samples were mixed with three biological replicates, and the number of replicates are presented in the figures. The standard deviation (SD) and P values were determined using Student’s *t*-test.

## Acknowledgments

We thank Zhangying Wang at Guangdong Academy of Agricultural Sciences and Hongbo Zhu at Guangdong Ocean University for providing plant materials, Guangyi Dai at South China Botanical Garden for providing technical support, and Xiaoyun Li at South China Normal University for providing the mScarlet and RUBY vectors. This work was supported by grants from the National Natural Science Foundation of China-Guangdong Natural Science Foundation Joint Project (U1701234), the National Natural Science Foundation of China (31970623), Guangdong Special Support Plan Project (2019TQ05N140), and the Guangzhou Municipal Science and Technology Project (202002030057).

## Author contributions

G.M., X.L. and X.H. designed the experiments. G.M., A.C., Y.W., S.L. and M.W. performed the experiments. G.M. and X.L. performed the data analysis. G.M., X.L. and X.H. wrote the manuscript.

