## Supplementary Figures for "A simple and efficient *in planta* transformation method based on the active regeneration capacity of plants"

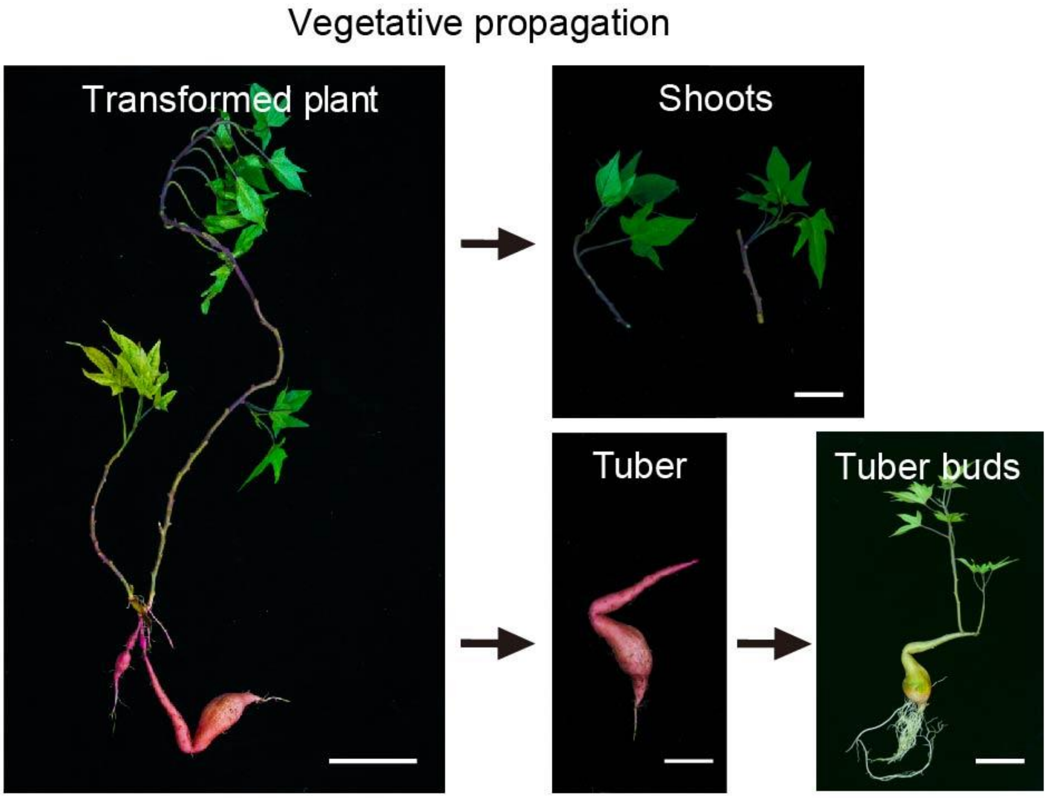


**Indentification of the transformed plants**

| Shoots | 1 | 2 | 3 | 4 | 5 | 6 | 7 | 8 | 9 | 10 |
|---|---|---|---|---|---|---|---|---|---|---|
| Positive lines | 0 | 0 | 0 | 0 | 0 | 0 | 1 | 2 | 0 | 0 |
| Total samples | 3 | 1 | 1 | 2 | 3 | 2 | 2 | 2 | 1 | 1 |

| Tuber buds | 1 | 2 | 3 | 4 | 5 | 6 | 7 | 8 | 9 | 10 | 11 | 12 |
|---|---|---|---|---|---|---|---|---|---|---|---|---|
| Positive lines | 0 | 0 | 0 | 0 | 0 | 0 | 0 | 2 | 1 | 3 | 0 | 0 |
| Total buds | 3 | 4 | 7 | 1 | 1 | 4 | 1 | 2 | 1 | 3 | 2 | 2 |
| Total samples | 10 plants | | | | | | | | | | | |

**Supplementary Fig. 1 Generation of transformed offspring of sweet potato**

The stem cuttings generated transgenic renascent leaves, lateral shoots, and tubers developed from adventitious roots. The table depicts the identification of these tissues by genotyping and GUS staining.

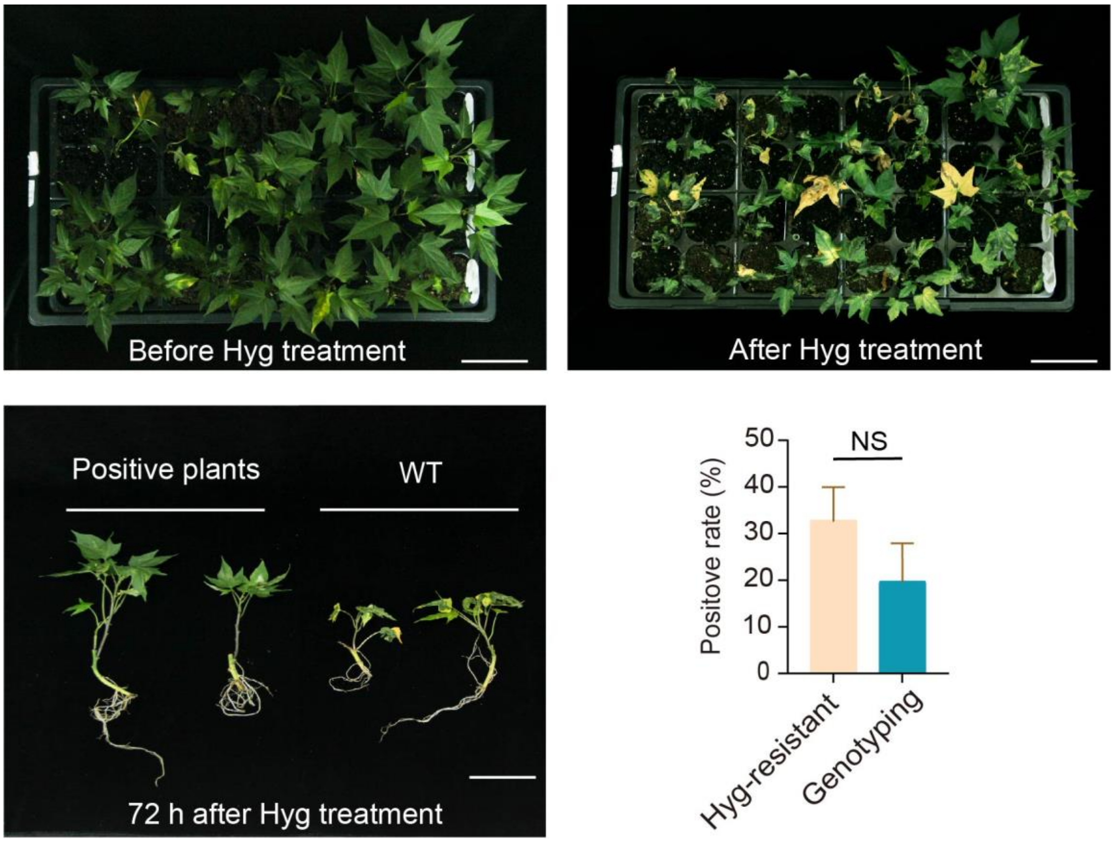


**Supplementary Fig. 2 Bulk selection of transgenic materials by hygromycin resistance**

Phenotypes of the injected plants before and after hygromycin (Hyg) treatment. Comparison of the morphology of the positive and WT plants after 72 h of applying Hyg. The bar chart depicts the positive rate of resistance and results of genotyping of screened plants. Positive rate = number of positive plants/total number of plants (%). The data are presented as the mean ± SD of ten biological replicates (two-tailed Student's *t*-test, NS, no significance, $P > 0.05$). Scale bar, 5 cm.

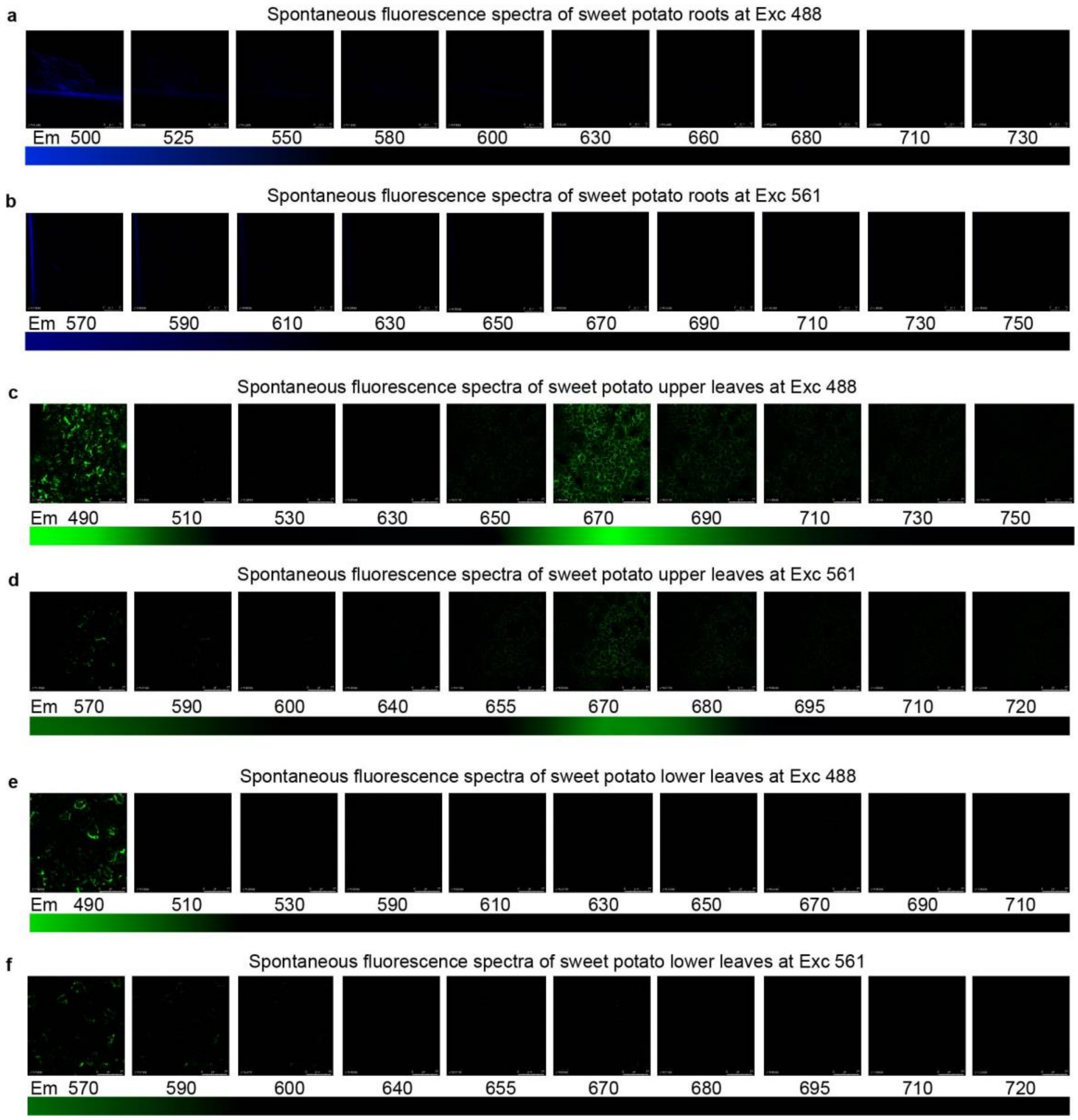


**Supplementary Fig. 3 Spontaneous fluorescence spectra of sweet potato tissues**

Spontaneous fluorescence spectra of the adventitious roots of sweet potato at **a**, Exc 488 nm and **b**, Exc 561 nm. Spontaneous fluorescence spectra of the upper leaves of sweet potato at **c**, Exc 488 nm and **d**, Exc 561 nm. Spontaneous fluorescence spectra of the lower leaves at **e**, Exc 488 nm and **f**, Exc 561 nm. Scale bar, 75 µm.

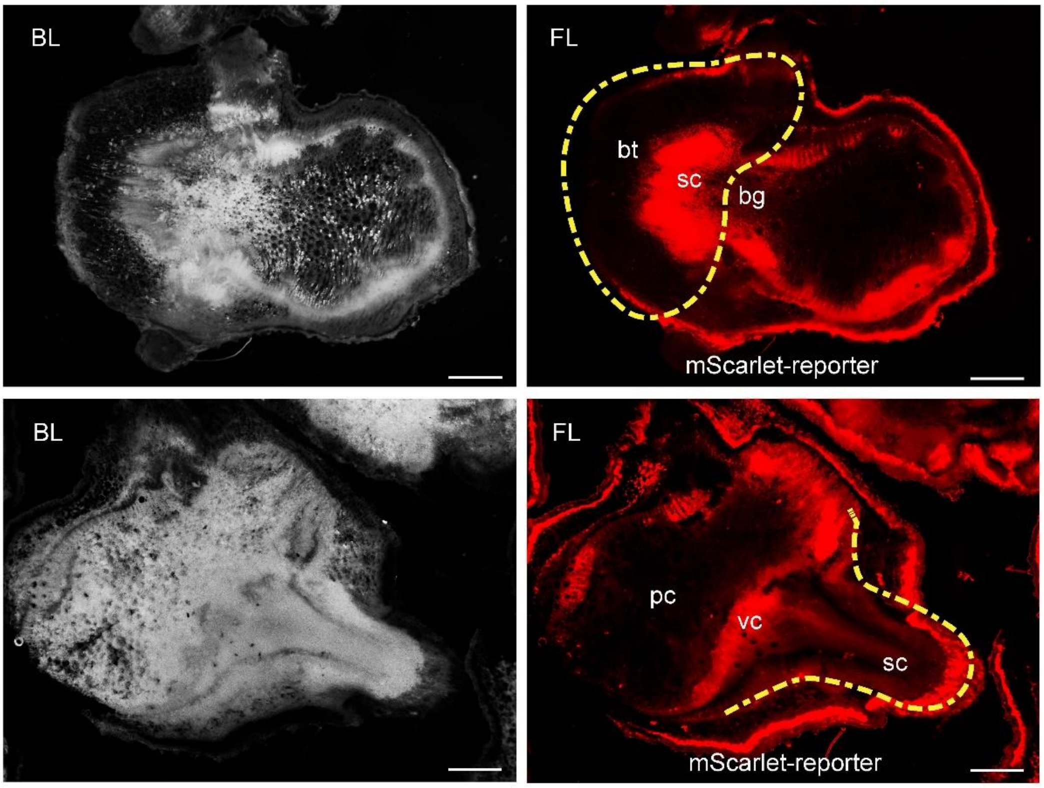


**Supplementary Fig. 4 Fluorescence signals due to mScarlet in the transfected renascence tissues**

The signals due to mScarlet in the cross-section of the lateral buds (upper) and adventitious roots (lower) of the transfected plants. The yellow line depicts the approximate area of the renascence tissues. The approximate spatial position of the lateral bud primordium was located between branch trace (bt) and branch gap (bg); stem cell, sc. The vascular cells (vc) are connected to the root primordium; parenchymal cell, pc; bright light, BL; fluorescent light, FL. Scale bar, 1 mm.

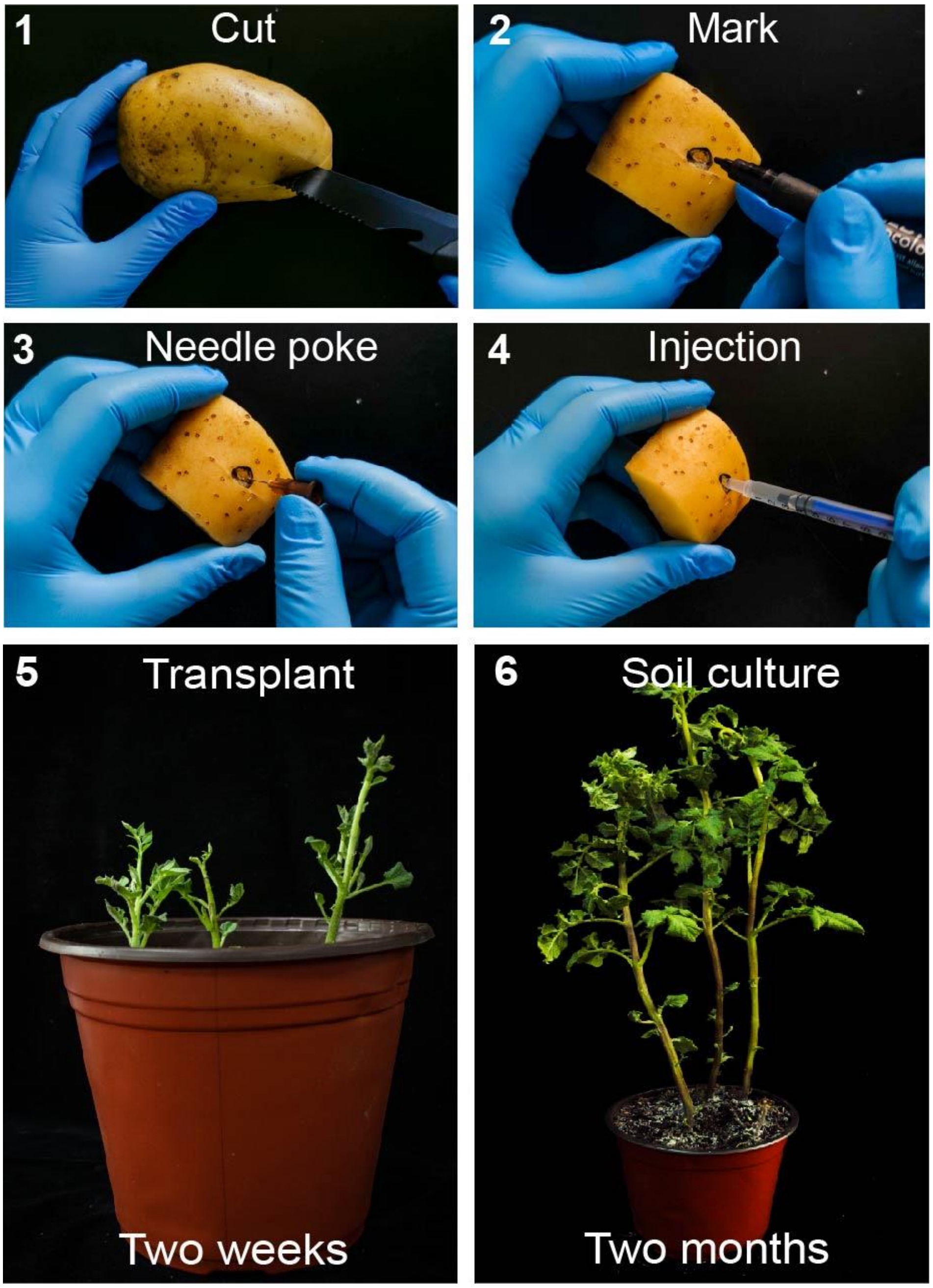


**Supplementary Fig. 5 Transformation procedure of potato by RAPID**

The fresh tubers of potato were cut and marked around the budding point. The tubers were poked with a needle, and the needle was inserted into the epidermis for injection. The transformed segments of tuber were transplanted in sandy soil. The new buds were unearthed after 1–2 weeks and subjected to further analysis for determining the positive transformants and phenotypic observations. The positive tubers were harvested after 6–8 weeks for subculture and mass propagation. Scale bar, 1 cm (up), 5 cm (left), 2 cm (right).
